# The economic value of vertebrate pollination in India

**DOI:** 10.1101/2024.12.20.629755

**Authors:** Ratheesh Kallivalappil, Sheena C. Cotter

## Abstract

The increasing urgency to understand the role and value of pollination in agriculture and plant conservation has elevated pollinators to a critical focus of research. While most crops—excluding cereals—are partially or entirely dependent on pollinators for fruit and seed production or benefit from improved yields due to their activities, the economic valuation of pollination services has primarily concentrated on insect pollinators globally and within India. However, comprehensive studies on the economic contributions of vertebrate pollinators in India remain absent. To address this gap, we evaluated the economic value of vertebrate pollination services for eight plant species by analyzing economic and production data from a published study. Using the methodologies of Klein et al. (2007), Gallai et al. (2009), and Gallai and Vaissiere (2009), we applied a contribution separation criterion to distinguish vertebrate pollination contributions from those of invertebrates. Our findings estimate the value of vertebrate pollination services at £2.86 million (D28.2 crore rupees), with bird pollinators contributing more than mammal pollinators. This study underscores the significant role of bat pollinators in supporting India’s economy and biodiversity through vital ecosystem services, highlighting their importance for conservation and policy initiatives.

## Introduction

Pollination is a vital ecosystem service, underpinning the functionality and sustainability of ecosystems (Costanza et al., 1997; Allen-Wardell et al., 1998; Kearns et al., 1998; Ricketts et al., 2008). Animal-mediated pollination facilitates the sexual reproduction of more than 90% of the estimated 250,000 species of angiosperms (Kearns et al., 1998). As a result, animals play a pivotal role in sustaining the diversity of flowering plants by supporting species richness, genetic variability, and the abundance of functional groups within ecosystems (Costanza et al., 1997; Balvanera et al., 2001; Díaz et al., 2006; Fontaine et al., 2005). The intricate interactions between plants and their pollinators are fundamental to maintaining the integrity and resilience of Earth’s biodiversity (Bawa, 1990; Biesmeijer et al., 2006; Heystek and Pauw, 2014).

Animal pollination is crucial for the sexual reproduction of most non-cereal crops and wild plants (McGregor, 1976; Williams, 1994; Nabhan and Buchmann, 1997; Kearns et al., 1998; Ashman et al., 2004). Approximately one-third of global food production relies on pollinator-dependent plants, including 86 of the 124 primary crops consumed worldwide (McGregor, 1976; Klein et al., 2007). In Europe, bees facilitate the pollination of 84% of cultivated crops (Williams, 1994), while biotic pollination enhances the quality or yield of 70% of 1,330 tropical crop species (Roubik, 1995). Pollinator-dependent crops also provide a significant share of essential nutrients in the human diet (Klein et al., 2007; Eilers et al., 2011).

Crop pollination has been recognized as an endangered ecosystem service (Allen-Wardell et al., 1998; Kearns et al., 1998; Kevan and Phillips, 2001; Potts et al., 2010). Pollinators significantly improve fruit quality and boost the economic value of crop production (Klatt et al., 2013; Garratt et al., 2014). However, their documented decline (Buchmann and Nabhan, 1996; Biesmeijer et al., 2006; Potts et al., 2010; Cameron et al., 2011) poses a risk to crop pollination, with serious economic repercussions and potential threats to global food security (Gallai et al., 2009).

Insects, including both managed and wild bees, are globally significant pollinators responsible for pollinating a diverse range of crops (McGregor, 1976; Free, 1993; Kremen et al., 2002; Greenleaf and Kremen, 2006; Winfree et al., 2007, 2008; Garibaldi et al., 2011, 2013). However, vertebrates also play a vital role in pollination, contributing to the reproduction of angiosperms and supporting biodiversity and ecosystem stability (Proctor et al., 1996). While research has primarily focused on invertebrates, vertebrate pollination—especially common in the tropics and less frequent in colder regions—remains underappreciated. Birds and mammals are effective pollinators of various cultivated plants, including economically important crops like dragon fruits (Hernandez and Salazar, 2012), durian and beans (Bumrungsri et al., 2008, 2009; Aziz et al., 2017), agave cacti (Kunz et al., 2011), and feijoa (Vogel et al., 1984; Stewart and Craig, 1989).

Quantifying the benefits of animal pollination is challenging due to its inherent complexities, such as incomplete data on both marketable and non-marketable products (Klein et al., 2007; Chaudhary and Chand, 2017). While numerous studies have assessed the economic value of pollination services (Losey and Vaughan, 2006; Klein et al., 2007; Gallai et al., 2009), the valuation process differs from that of plants. A plant’s economic value is typically calculated by multiplying production quantity by its annual market rate, whereas the value of pollination services involves multiplying the economic value of annual plant production by its pollinator dependence rate, which reflects how reliant a plant is on pollinators for reproduction (Klein et al., 2007; Gallai et al., 2009; Gallai and Vaissiere, 2009).

In India, research on pollination services has predominantly focused on insect pollination (Goyal, 1993; Chaudhary, 1998; Chaudhary and Chand, 2017), with little attention to vertebrate pollinators. As a result, the contribution of vertebrate pollinators to the Indian economy remains largely unknown. For example, the ecological services provided by fruit bats—key pollinators and seed dispersers for many tropical plants (Cox et al., 1991; Fujita and Tuttle, 1991)—are undervalued by the public and conservation planners (Singaravelan et al., 2009). Traditionally, bats have been hunted for food and medicine and persecuted as crop pests, partly due to their outdated “pest” designation under the Indian Wildlife Protection Act of 1972, which was only recently revoked.

The absence of quantitative data on vertebrate pollinators, importantly fruit bats’, ecological contributions hamper their recognition and protection in conservation programmes. Therefore, we attempt to estimate their contribution to the Indian biodiversity and economy for effective conservation.

## Materials and Methods

### Extracting production and economic value data

To assess the economic value of selected plant species, we collected data on annual economic value (range price per kilogram in Indian Rupees) and total annual production of plant parts (flowers, fruits, seeds, stem-bark, root, leaf) in metric tons (MT) were extracted from the study “Medicinal plants in India: An assessment of their demand and supply” conducted by Goraya and Ved (2017) under the National Medicinal Plants Board and Indian Council of Forest Research Education, Government of India. The annual economic value of each plant species was determined by summing the market value of its relevant plant parts used within a year. Our survey using other databases (for examples, National Medicinal Plant Board, Department of Agriculture and Farmers Welfare, and the Ministry of Commerce and Industry - Government of India) did not find sufficient data to analysis the pollination service value for the plant species that have been recorded as vertebrate pollinated plants by Kallivalappil et al. (2024). Therefore, we analyse this for the plant species that has adequate data (Table 2).

### Categorising vertebrate-pollinated plants

We categorized all plants into self-incompatible and self-compatible groups based on their reproductive systems. Self-incompatible plants, which rely exclusively on foreign pollen for reproduction, were classified as “essential,” assuming no seed or fruit production would occur without pollinator activity. For self-compatible plants, capable of reproducing with self-pollen, we considered the contribution of pollinators to seed/fruit production. We assumed that higher pollinator activity leads to increased production rates (Klein et al., 2007), as self-pollination typically results in minimal yields for most plants. The percentage of seed/fruit production for each species was derived from existing studies that reported results from pollination experiments, including manual pollination or natural pollinator visits to flowers.

### Assigning Dependency Rate

Subsequently, we assigned a dependency rate to each plant category, reflecting the degree to which plants rely on pollinators for their production (Klein et al., 2007; Gallai et al., 2009; Gallai and Vaissiere, 2009). This classification helps quantify the extent of pollinator contributions to plant reproduction and production, distinguishing whether a plant species is entirely, partially, or minimally dependent on animal pollinators. The plants were categorized into six groups, each assigned a *Dependency Rate* based on their reliance on pollinators for seed or fruit production (Table 1).

**Table 1:** Plant species grouped based on their degree of dependency for pollinators and the *Dependency Rate* of each group according to Klein et al. (2007) Gallai et al. (2009) and Gallai and Vaissiere (2009).

| Groups | % of reduction without pollinators | Dependency Rate<br>(DR) |
| --- | --- | --- |
| Essential | Pollinators essential (production reduction by $\geq 90\%$) | 0.95 |
| Great | Great production increase/ animal pollinators are strongly needed (40 - < 90% reduction) | 0.65 |
| Modest | Modest production increase/animal pollinators are clearly beneficial (10 - <40% reduction) | 0.25 |
| Little | Little production increase /some evidence suggests that animal pollinators are beneficial (>0 - <10% reduction) | 0.05 |
| No increase | No production increase with animal-mediated pollination | 0 |
| Unknown | Empirical/ real studies are missing | 0 |

#### Estimating Economic value of animal pollination

We estimated the economic value of animal pollination services (including both vertebrates and invertebrates) using the methodology outlined by Chaudhary and Chand (2017). In some cases, a single plant species was pollinated by both groups, making it necessary to assess their combined contribution. To determine this, we calculated the mean value of total annual production (in metric tons) and annual production values for each plant species. The production values were then converted to kilograms. Finally, the economic value for each plant species was derived by multiplying the annual production with its corresponding annual market rate.

EV = QT x AR

Where EV = Economic value of plant products annually

QT = Production Quantity of plants annually

AR = Annual Rate of plant products

Thus, the economic value of pollination services (EVP) for each plant species was calculated by multiplying the economic value (EV) of the annual plant production by the *dependency rate* (Table. 1).

EVP = EV x DR

DR = Dependence Rate

#### Economic value of vertebrate pollination services

We found certain plant species were pollinated by both invertebrates and vertebrates (Table 2), necessitating the separation of invertebrate contributions to determine the relative contribution of vertebrates. Due to the lack of studies estimating the individual contributions of invertebrates and vertebrates, we calculated the total economic value of animal pollination and divided it by the number of recorded pollinators (both invertebrate and vertebrate) for each plant species. This approach assumes equal contribution of each pollinator species to plant reproduction. While this assumption may not hold—abundant species, for example, likely contribute more significantly to production success—these nuances could not be accounted for due to unavailable data. Indian studies predominantly focus on identifying pollinators or breeding systems rather than quantifying individual pollinator contributions.

**Table 2:**
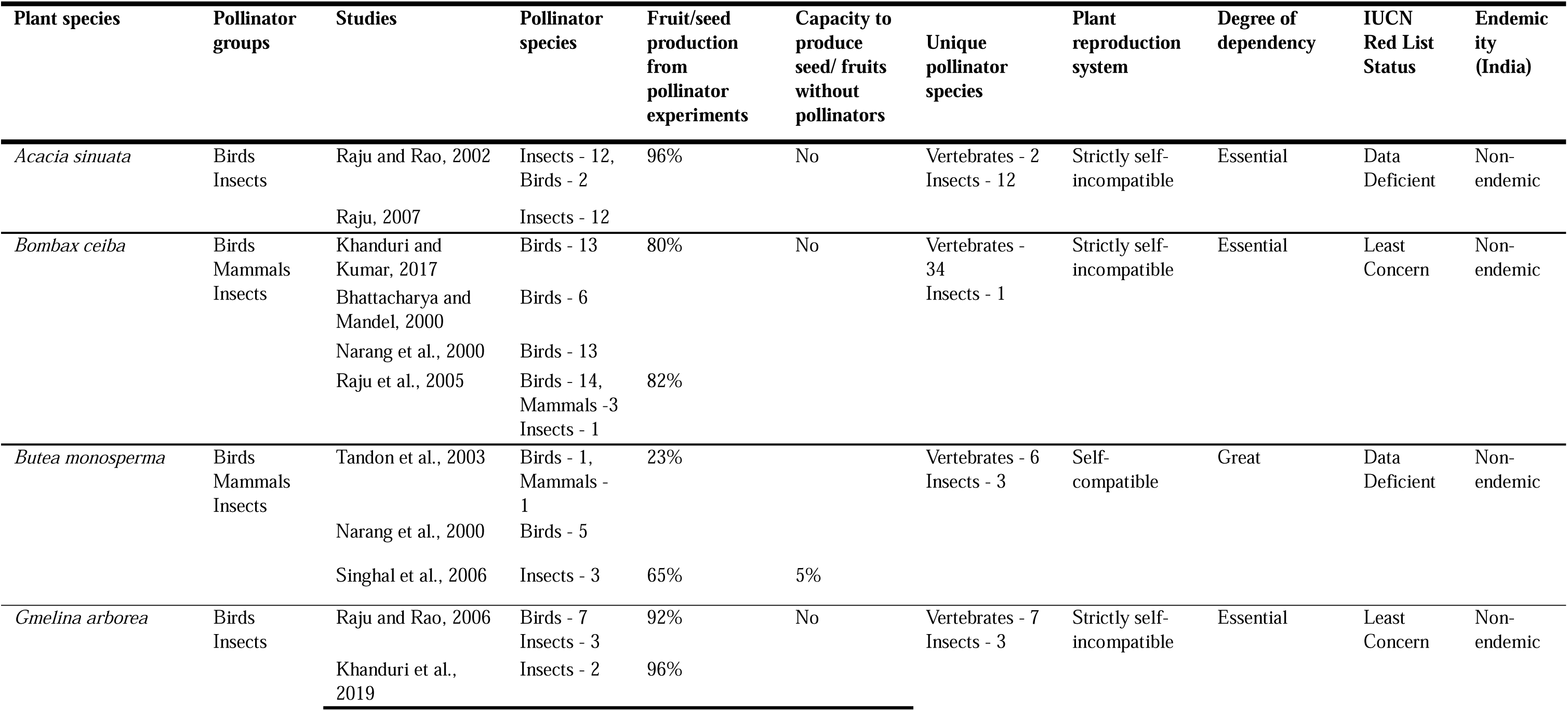
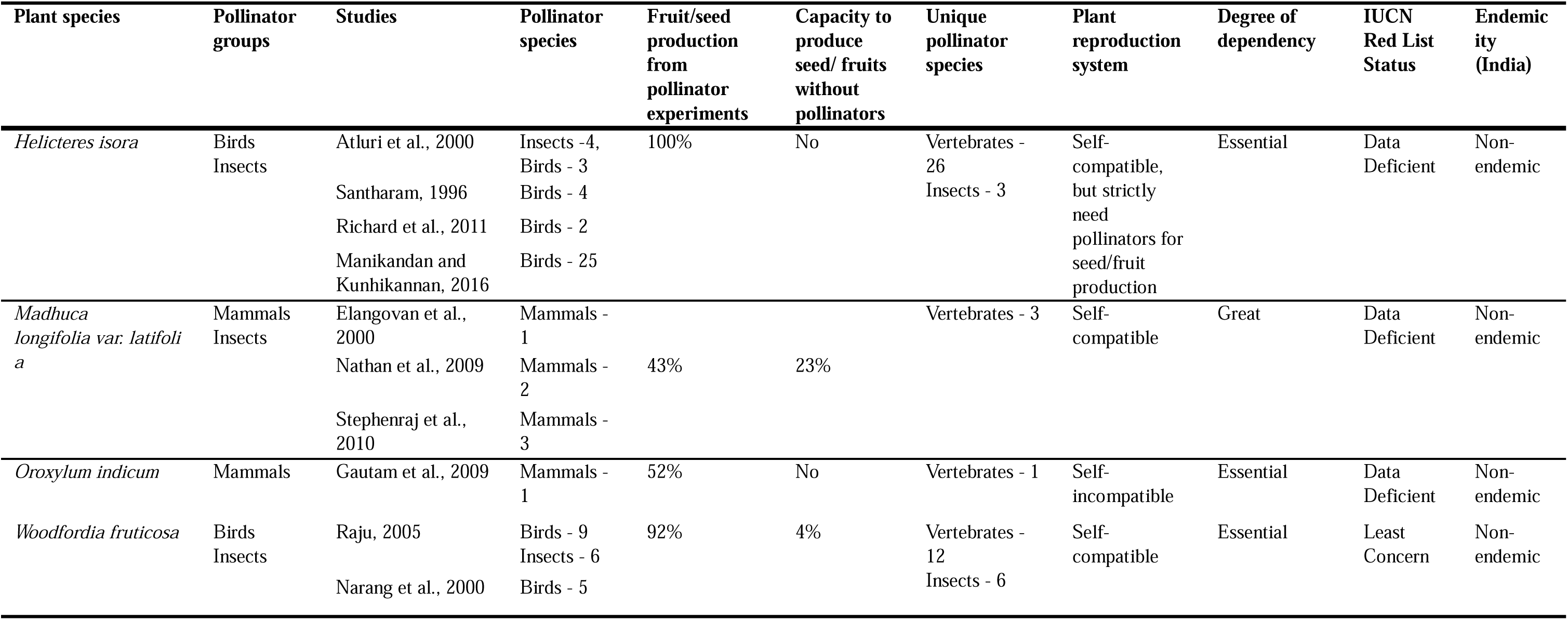
Plant species used for the assessment of economic value of vertebrate pollination services. Each plant species has been categorised into essential, great, modest, little, no increase and unknown based on the degree of dependency for pollinators. The degree of plant’s dependency was based on the reproductive system (for self-incompatible) and fruit/seed production rate (for self-compatible) of plant species as per the pollination experiment studies. The value of capacity to produce seed/fruits without pollinators are derived from the respective studies that cited for each plant species.

Therefore, we isolated the vertebrate pollinators’ contribution by either assessing the combined contribution of all vertebrate pollinators for each plant species or subtracting the invertebrate contribution from the total value of animal pollination services.

Contribution of Individual Pollinator = Value of animal pollination/total number of pollinators

Economic Value of Vertebrate Pollination Services = Sum of total vertebrate pollinators

Additionally, we explored the contribution of individual vertebrate pollinators’ service value to each plant species by dividing the pollination value of each plant by its total pollinator species.

Pollination service value of each vertebrate = Total vertebrate pollination value / Total vertebrate pollinator species

## Results

Our study showed that the degree of dependency for pollination varied across the plant species (Table 2). The economic value of animal pollination services for the assessed 8 plant species was, on average, £4,614,604 or £4.6 million (D455,000,000 or D45.5 crores; Crore is an Indian system of units. One crore is equal to 10 million, Table 3). However, the total economic value of vertebrate pollination services was estimated as £2,856,115 or £2.8 million (D281,612,946 or D28.2 crores; Table 3).

**Table 3:** Assessment of the economic value of vertebrate pollination services. The abbreviations used in the table, MT (Metric ton), Rs (Indian Rupees), Kg (Kilogram), EV (Economic Value), EVP (Economic Value of Pollination). The average exchange rate of rupees 98.6/pound in 2014 was used to convert to pound values. The production and economic value data have been taken from Goraya and Ved, 2017.

| Plants | Pollinator Groups | Production (MT) / Year | Average production (MT) | Average production in Kg / Year | Rate (Rs/Kg) | Average Rate (Rs/kg) | EV (Rs) | Animal Pollination | Dependency rate | EVP animal (Rs) | Total pollinator species | Individual pollinator's contribution | Total vertebrate pollinators | EVP Vertebrate (in Indian Rupees) | EVP Vertebrate (in Pounds Sterling) |
| --- | --- | --- | --- | --- | --- | --- | --- | --- | --- | --- | --- | --- | --- | --- | --- |
| *Acacia sinuata* | Bird, Insect | 1,000 - 2,000 | 1,500 | 15,00,000 | 25 - 95 | 60 | 90,000,000 | Essential | 0.95 | 85,500,000 | 14 | 6,107,142 | 2 | 12,214,285 | 123,877 |
| *Bombax ceiba* | Bird, Mammal, Insect | 100 - 200 | 150 | 150,000 | 140-160 | 150 | 22,500,000 | Essential | 0.95 | 21,375,000 | 35 | 610,714 | 34 | 20,764,285 | 210,591 |
| *Butea monosperma* | Bird, mammal, Insect | 200 - 500 | 350 | 350,000 | 150-350 | 250 | 87,500,000 | Great | 0.65 | 56,875,000 | 9 | 6,319,444 | 6 | 37,916,666 | 384,550 |
| *Gmelina arborea* | Bird, Insect | 500 - 1,000 | 750 | 750,000 | 35-40 | 37.5 | 28,125,000 | Essential | 0.95 | 26,718,750 | 10 | 2,671,875 | 7 | 18,703,125 | 189,687 |
| *Helicteres isora* | Bird, Insect | 100 - 200 | 150 | 150,000 | 20-30 | 25 | 3,750,000 | Essential | 0.95 | 3,562,500 | 30 | 118,750 | 26 | 3,087,500 | 31,313 |
| *Madhuca longifolia* var. *latifolia* | Mammal | 200 - 500 | 350 | 350,000 | 75-100 | 87.5 | 30,625,000 | Great | 0.65 | 19,906,250 | 3 | 6,635,416 | 3 | 19,906,250 | 201,889 |
| *Oroxylum indicum* | Mammal | 500 - 1,000 | 750 | 750,000 | 30-40 | 35 | 26,250,000 | Essential | 0.95 | 24,937,500 | 1 | 24,937,500 | 1 | 24,937,500 | 252,916 |
| *Woodfordia fruticosa* | Bird, Insect | 2000 - 5,000 | 3,500 | 3,500,000 | 60-70 | 65 | 227,500,000 | Essential | 0.95 | 216,125,000 | 18 | 12,006,944 | 12 | 144,083,333 | 1,461,291 |
| **Total** |  |  | **7,500** | **7,500,000** |  |  | **516,250,000** |  |  | **455,000,000** |  |  |  | **281,612,946** | **2,856,115** |

### Economic value of bird and mammal pollination services

The economic value of bird pollination services was, on average, £2,312,443 or £2.3 million (LJ228,006,894 or LJ22.8 crores, Table 4). This was 81% of the total vertebrate pollination service values. This amount was the collective contribution of 49 bird species who involved in the pollination of assessed plants species (Table 4). The purple sunbird (*Cinnyris asiaticus*) provided the highest pollination service value of £282,300 (LJ27,834,871 or LJ2.8 crores, N = 6 plants). Similarly, the purple-rumped sunbird (*Leptocoma zeylonica*) provided the pollination value of £212,015 (LJ20,904,712 or LJ2.1 crores, N = 4 plants, Table 4). These two bird species together contributed a pollination value of £494,316 (LJ48,739,583 or LJ4.9 crores), which was 21% of the total bird pollination values (Table 4).

**Table 4:**
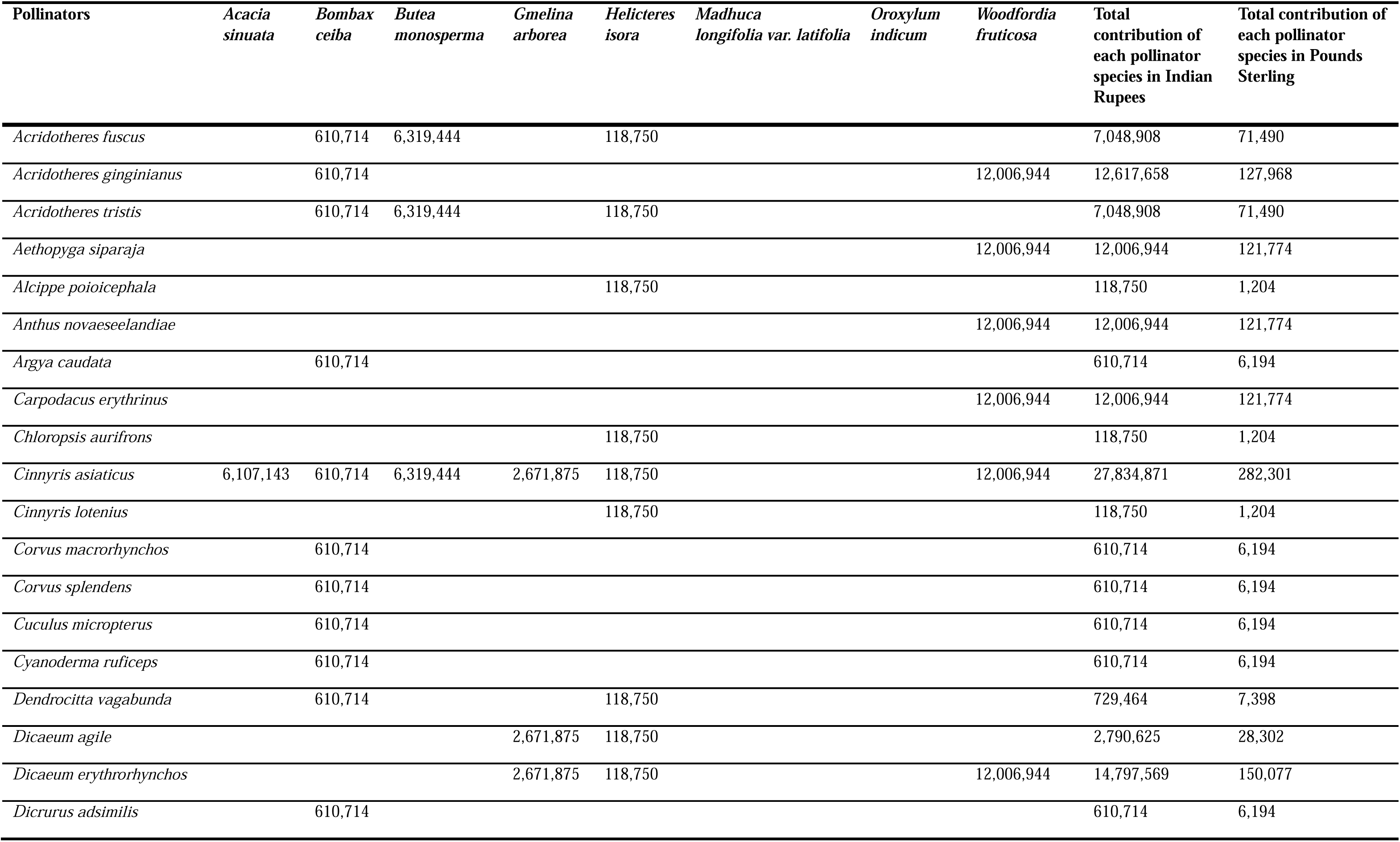
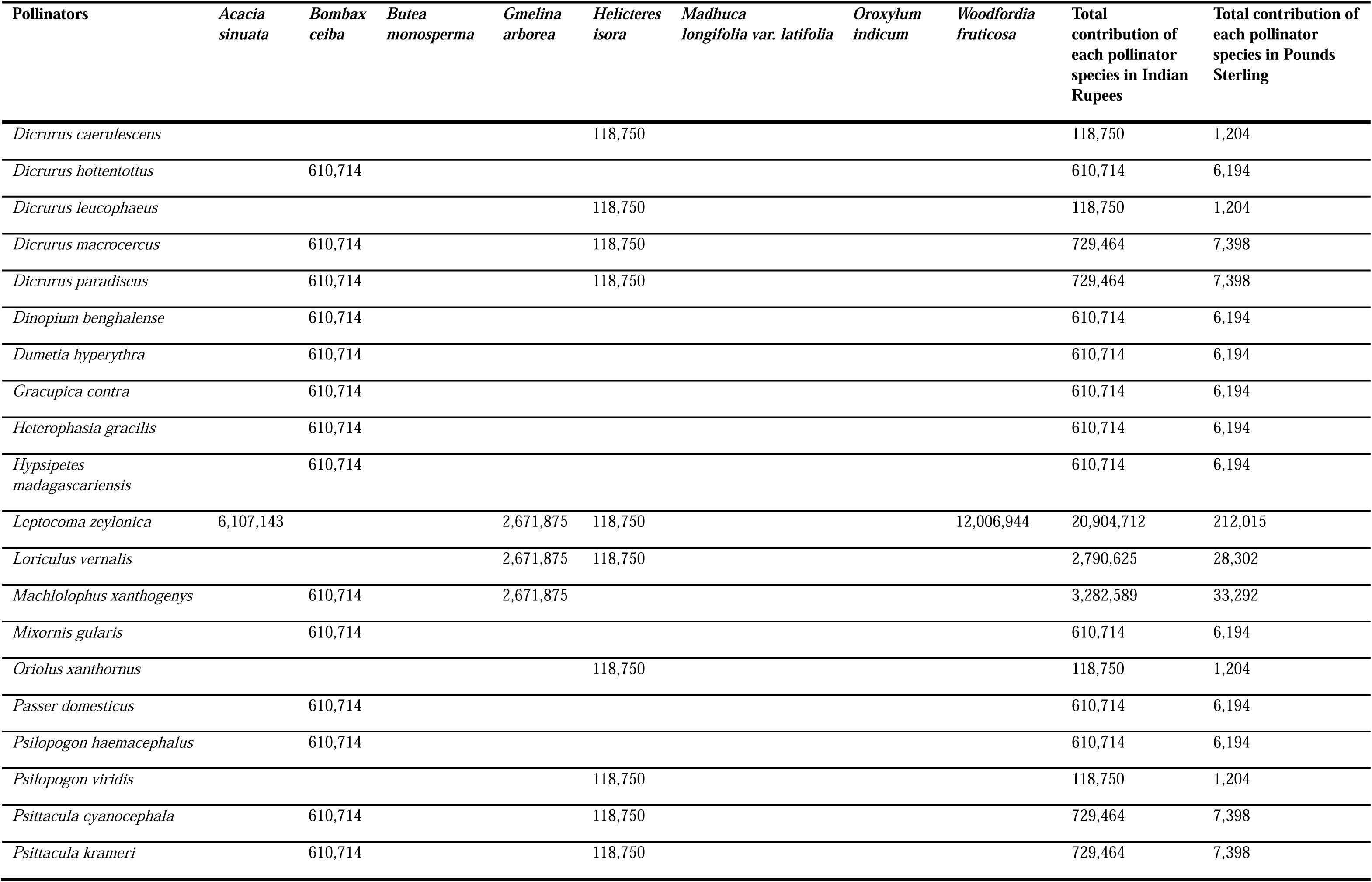
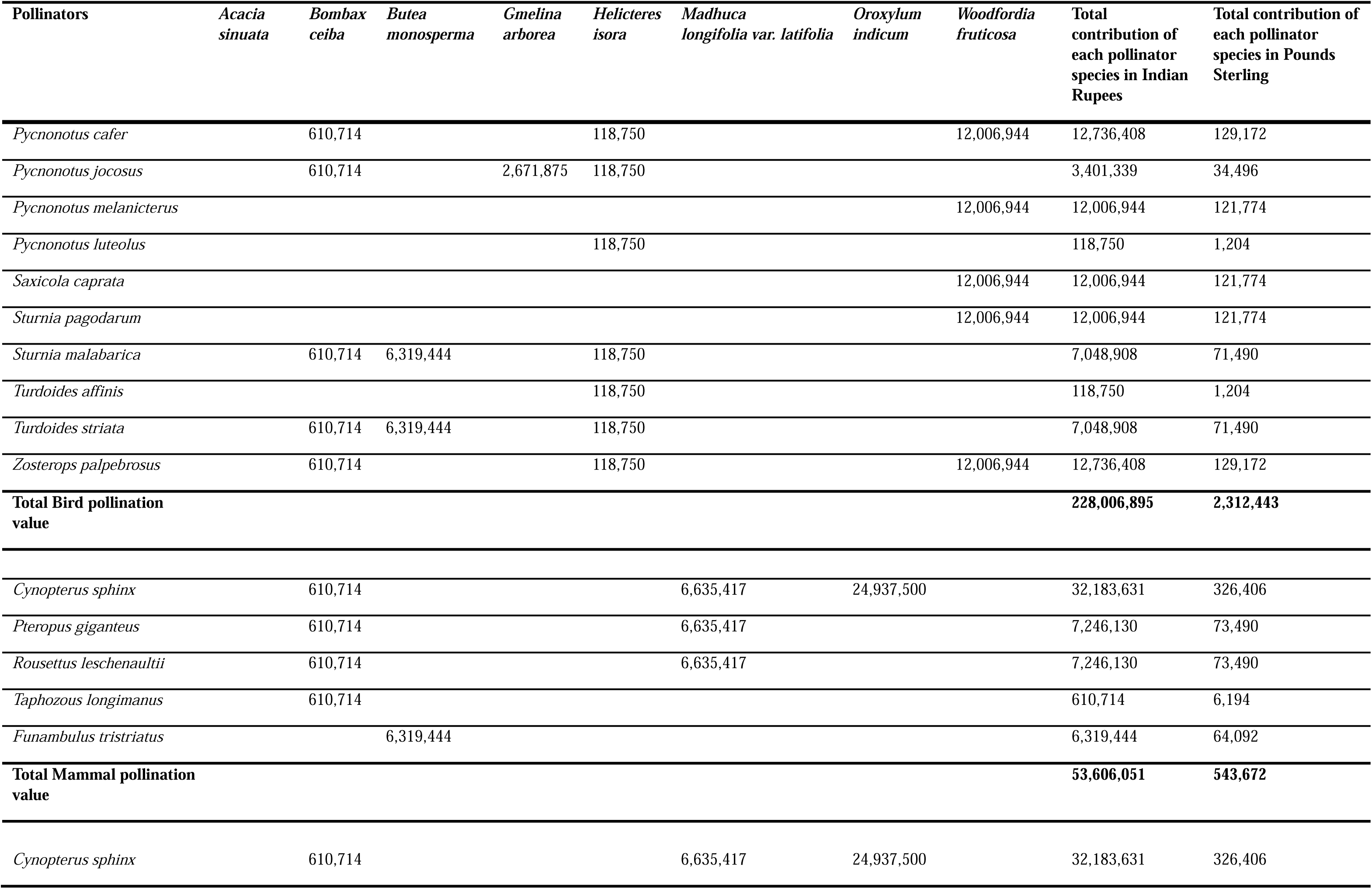
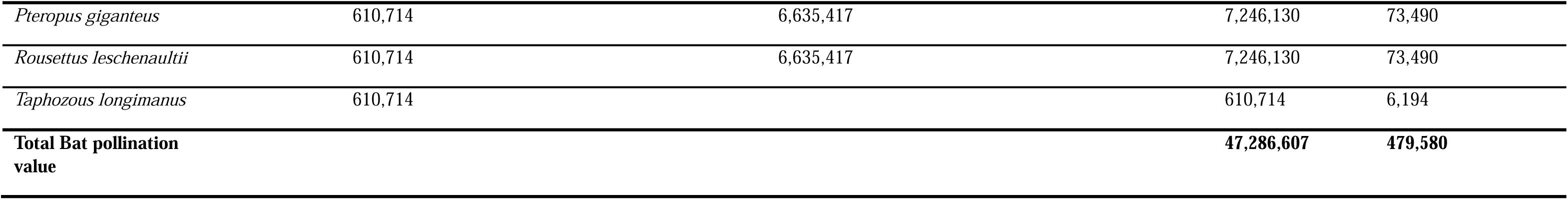
Assessment of economic contribution of each vertebrate pollinator species on specific plant species.

Consequently, the total economic value of mammal pollination services was, on average, £543,672 (LJ53,606,051 or LJ5.4 crores, Table 4). It was 19% of the total vertebrate pollination values. Among this, the pollination service value of bats was £479,580 (LJ47,286,607 or 4.7 crores), that was 88% of the total mammal pollination values (Table 4). The Greater short-nosed fruit bat (*Cynopterus sphinx*) contributed the highest value among all the mammal pollinators £326,405 (LJ32,183,630 or 3.2 crores, N = 3 plants) and it was 60% of the total mammal pollination values (Table 4).

## Discussion

Numerous studies worldwide have assessed the economic value of pollination services, with a predominant focus on insect pollinators (e.g., Klein et al., 2007; Gallai et al., 2009; Gallai and Vaissiere, 2009). In India, such studies have primarily concentrated on insect pollinators and crop species, leaving the contribution of vertebrate pollinators largely overlooked. Utilizing available economic and production data for 8 out of 89 vertebrate-pollinated plant species, we estimated the economic value of vertebrate pollination services in India to be approximately £2.8 million (LJ281,612,946 or LJ28.2 crores). The actual total value of vertebrate pollination services is likely to be significantly higher.

This underestimation stems from the limited focus on vertebrate-pollinated plants, potentially leading to an incomplete understanding of their contributions. Vertebrates may pollinate many more economically valuable plant species in India than currently documented. This information gap not only hinders the conservation of these pollinators and their associated plants but also impacts the livelihoods of dependent communities and national revenue.

In 2014-15, India’s commercial demand for herbal raw drugs was estimated at 5,12,000 metric tons, with exports of 1,34,500 metric tons (Goraya and Ved, 2017). Several of these medicinal plants are pollinated by birds and mammals. Beyond medicinal applications, these plants serve additional purposes, including use as fodder, timber, food, fibre, and fuel. The lack of comprehensive data restricts our ability to fully evaluate the extensive economic impact of vertebrate pollination services.

The value of bird pollination services was (£2.3 million) higher than the mammal pollination services (£543,671). Birds are globally recognized for their vital ecosystem services (Whelan et al., 2008) and contribute to the pollination of numerous economically important plant species (Vogel et al., 1984; Stewart and Craig, 1989; Nabhan and Buchmann, 1997). However, studies quantifying the economic value of bird pollination remain sparse, highlighting the need for further research. While bird pollinators likely contribute a small proportion of the 35% global food production dependent on animal pollination (Klein et al., 2007), their role is nonetheless critical.

In comparison to birds, data on the economic value of pollination services by bats is even more limited. In Peninsular Malaysia, annual sales of bat-pollinated petai (*Parkia speciosa* and *Parkia javanica*) exceed £430,000 (more than US$1 million; Ng, 1980). In Indonesia, bat-pollinated crops such as petai and duku (*Lansium domesticum*) generate annual revenues exceeding £2,260,000 (more than US$4 million; Fujita and Tuttle, 1991). Durian pollination services by bats in Indonesia have been valued at approximately £88/ha/fruiting season (US$117/ha), amounting to £337,000 (US$450,000) in a single village (Ober and Tsang, 2019). Similarly, in Mexico, the export value of the bat-pollinated *Agave tequilana*, used for tequila production, was an astounding £785 million (US$1.2 billion) in August 2015 (Trejo-Salazar et al., 2016). The ongoing decline of pollinator and plant populations poses significant threats to biodiversity and the economy, often leading to an underappreciation of the essential benefits these pollinators provide to humanity.

## Conclusion

This study estimated the pollination service value of vertebrate pollinators using 8 plant species. The study highlights the importance of birds and bats as important ecosystem service providers. Historically, the absence of quantitative data on these services and lenient regulations have hindered the protection of fruit bats in India, contributing to significant declines in their populations and the ecosystem services they support. To address this, we advocate for the implementation of targeted conservation strategies for bat pollinators through the systematic documentation of natural caves, strict prohibitions on human intrusion into these habitats, and community-led programmes to monitor and protect roosts and foraging sites near human settlements.

## Supporting information

https://zenodo.org/records/10926527

## Acknowledgements

We thank Goraya and Ved for the extensive study on the economic value of medicinal plants in India and making the data available along with appreciating the Ministry of Social Justice and Empowerment, Government of India, for the financial support to RK through the National Overseas Scholarship. We also thank the University of Lincoln for providing facilities and technical support.

## Author contributions

The study was conceptualised and designed by both Ratheesh Kallivalappil and Sheena C. Cotter. The literature survey was conducted by RK, and the first draft of the manuscript was written and edited by both RK and SC. Both authors read and approved the manuscript.

## Data Availability

This study did not generate any quantitative datasets.

